# Beyond cortical geometry: brain dynamics shaped by rare long-range connections

**DOI:** 10.1101/2024.04.09.588757

**Authors:** Jakub Vohryzek, Yonatan Sanz-Perl, Morten L. Kringelbach, Gustavo Deco

**Author notes:** Jakub Vohryzek **Email:**. **Author Contributions:** J.V., G.D. and M.L.K. designed research; J.V., Y.S-P., M.L.K. and G.D. performed research; J.V., Y.S-P. and G.D. contributed new reagents/analytic tools; G.D., J.V. and Y.S-P. analyzed data; J.V. G.D. and M.L.K. wrote the paper.

## Abstract

A fundamental topological principle is that the container always shapes the content. In neuroscience, this translates into how the brain anatomy shapes brain dynamics. From neuroanatomy, the topology of the mammalian brain can be approximated by local connectivity, accurately described by an exponential distance rule (EDR). The compact, folded geometry of the cortex is shaped by this local connectivity and the geometric harmonic modes can reconstruct much of the functional dynamics. However, this ignores the fundamental role of the rare long-range cortical connections, crucial for improving information processing in the mammalian brain, but not captured by local cortical folding and geometry. Here we show the superiority of harmonic modes combining rare long-range connectivity with EDR (EDR+LR) in capturing functional dynamics (specifically long-range functional connectivity and task-evoked brain activity) compared to geometry and EDR representations. Importantly, the orchestration of dynamics is carried out by a more efficient manifold made up of a low number of fundamental EDR+LR modes. Our results show the importance of rare long-range connectivity for capturing the complexity of functional brain activity through a low-dimensional manifold shaped by fundamental EDR+LR modes.

**Significance Statement:** Explaining how structure of the brain gives rise to its emerging dynamics is a primary pursuit in neuroscience. We describe a fundamental anatomical constraint that emphasises the key role of rare long-range connections in explaining functional organisation of the brain in terms of spontaneous and task-evoked activity. Specifically, this constraint unifies brain geometry and local connectivity through the Exponential Distance Rule while considering the long-range exceptions to this local connectivity as derived from the structural connectome. In addition, when using this structural information, we show that the task-evoked brain activity is described by a low-dimensional manifold of several modes suggesting that less is more for the efficient information processing in the brain.

## Introduction

How brain underlying anatomy shapes functional dynamics is an unresolved question being studied from the perspective of network neuroscience (1), brain modelling (2), graph signal theory (3) and neural field theories with different assumptions on the underlying anatomy (4, 5). Therefore, the choice of underlying anatomical features is of paramount importance in deriving the most simple and parsimonious description of the emerging spatiotemporal brain dynamics.

In previous work on retrograde tract tracing in non-human primates, Kennedy and colleagues have shown that the brain white-matter wiring can be analytically approximated by the Exponential Distance Rule (EDR) (6). This rule explains the local connectivity of the brain solely in terms of the geodesic distance between points on the cortical surface. And so, it follows that the compact, folded geometry of the cortex with its many sulci and gyri is formed by this local connectivity. This corollary implies that the brain anatomical wiring and cortical geometry are the two sides of the same coin, and it makes sense to speak of them in agreement. Furthermore, this reflects theoretical work showing that the heat kernel (exponential) is the optimal solution for minimising distance between neighbouring points (7). Indeed, recent work has suggested that the cortical geometry alone (as a proxy for the underlying anatomical connectivity) can be considered as an important feature driving brain spatiotemporal activity (5, 8, 9).

However, after deriving the EDR Henry Kennedy famously said; “I am not interested in the EDR itself but mainly the exceptions to the rule”. Indeed, Kennedy and colleagues have shown that in addition to the EDR, the brain possesses a small subset of rare long-range (LR) exceptions to the EDR of brain wiring (10, 11). Furthermore, new evidence using turbulence has demonstrated the fundamental role of the rare long-range anatomical connections in shaping optimal brain information processing (12). Intuitively, brain cortical foldings defined according to the EDR are indeed the optimal way for brain wiring but they don’t reflect the rare long-range connections i.e. it is for example impossible to fold anterior-posterior brain regions in a meaningful way. Therefore, we suggest that the unique contribution of these rare long-range cortical connections’ changes disproportionately the topological structure of the brain wiring in such a way as to optimise the information processing of the brain. In this work we test this hypothesis that EDR and LR exceptions are fundamental to the parsimonious description of the emerging spatiotemporal dynamics.

In the natural world, a fundamental principle that governs the dynamics of a system constrained by its structure in numerous physical and biological phenomena is the mathematical framework of harmonic modes. Standing wave patterns manifest in many context such as in music with sound-induced vibrations of a guitar string, in physics with the electron wave function of a free particle described by the time-independent Schrödinger equation, or biology with patterns emerging within complex dynamical systems like reaction-diffusion model (13). The beauty of the mathematical formalism of this phenomenon is that it links in a single equation, the Helmholtz equation, the specific structure on which the spatiotemporal pattern emerges together with the temporal description in terms of oscillations and spatial description in terms of patterns of synchrony of the standing wave pattern itself.

Here we used Laplacian decomposition of four different graph representations of the underlying anatomy to derive anatomical brain modes: exponential-distance rule (EDR) (6) and long-range exceptions (EDR+LR), geometry-based modes (geometry) and EDR modes (EDR binary and EDR continuous) (**Figure 1A, 1B, 1C**). Our results show that EDR+LR achieves statistically better reconstruction of long-range functional connectivity (FC) compared to the other mode representations (**Figure 1D**). Furthermore, pertinent to time-critical information processing, we show that a small subset of modes achieves a disproportionately high reconstruction of task MRI activity. When this subset of modes is considered, EDR+LR achieves better reconstruction for the 47 HCP tasks compared to the geometric mode representations, suggesting that less is more for information processing in the brain (**Figure 1E**).

**Figure 1.**
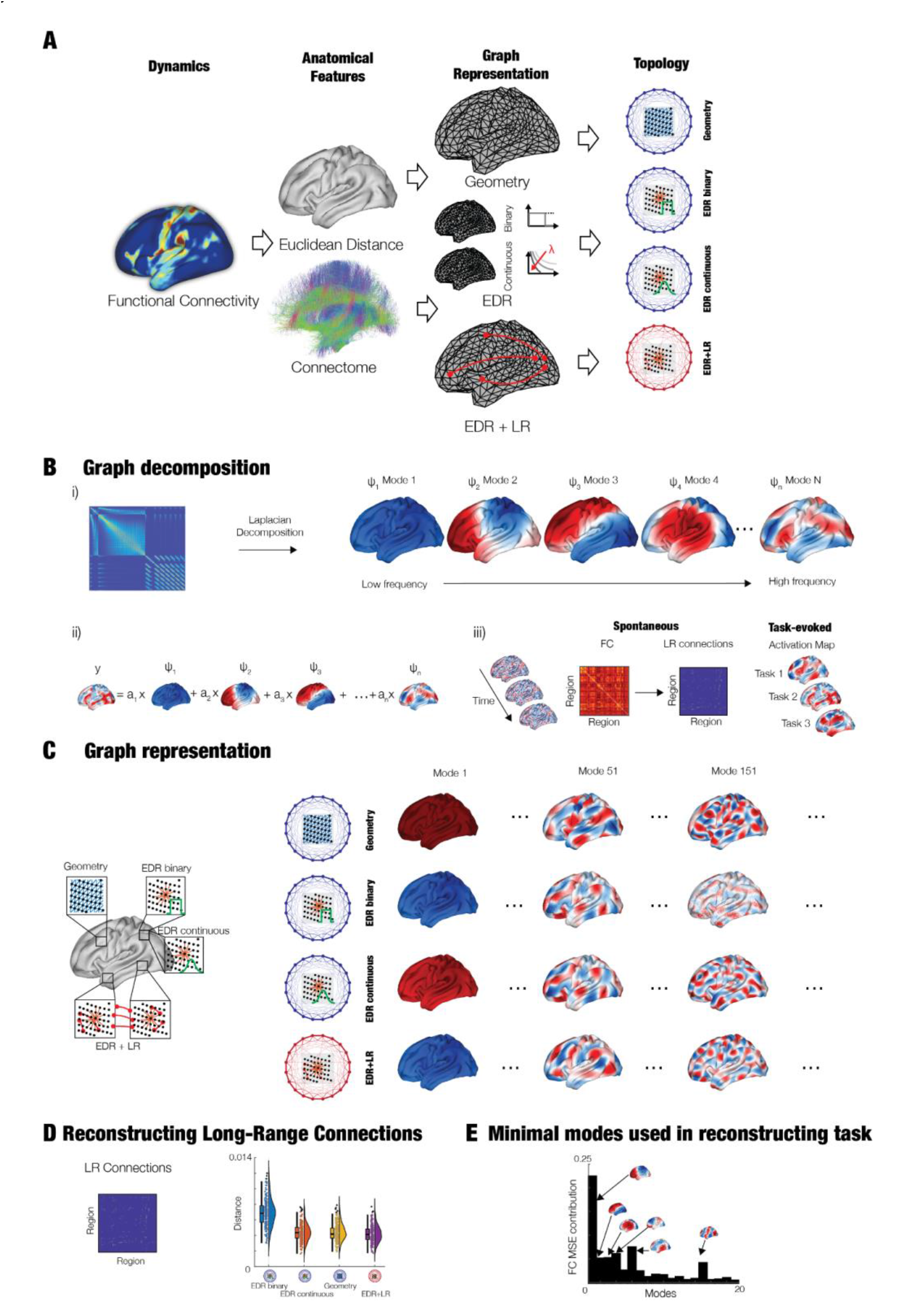
The crucial role of long-range connectivity for accurately describing whole-brain dynamics. **A)** The functional dynamics measured with fMRI emerge from the underlying anatomical structural connectivity which can be represented as graphs. Here, we study the four main graph representations: 1) geometrical modes (5); 2) exponential distance rule (EDR, binarized); 3) EDR (continuous) and 4) EDR with long-range exceptions (EDR+LR). **B)** With regards to the graph representations, i) the different modes are derived from applying the Laplace decomposition on the graph representation by solving the eigenvalue problem. The different modes are in ascending spatial frequency. ii) These modes are used to reconstruct the fMRI activity by a linear combination of their contributions. iii) This is used to reconstruct the spontaneous fMRI activity, and particularly the functional long-range connectivity exceptions (derived as high-correlation values, >0.5 correlation, and over a long Euclidean distance, >40mm, see *Methods*), as well as all the 47 task fMRI activation maps. **C)** The four different graph representations were constructed and decomposed into their associated modes. **D)** Demonstrating the importance of long-range connections, EDR+LR achieves a superior reconstruction of long-range fMRI connectivity compared to geometric, EDR (binary) and EDR (continuous) graph representations. **E)** Equally important, the EDR+LR needs fewer modes to reconstruct task data compared to the three other graph representations, demonstrating the importance of long-range connectivity. Parts of the figure have been modified from work by Pang and colleagues (5).

## Results

### EDR+LR reconstructs FC-SC long-range connectivity

To examine how exponential distance rule with long-range exceptions can describe brain activity, we derived the EDR+LR harmonic modes from the EDR matrix fitted to the structural connectome with lambda of 0.162 and added the long-range exceptions to the EDR defined in terms of three standard deviations from a given Euclidean distance range larger than 40mm. We constructed the normalised graph Laplacian and solved its eigenvalue problem (**Figure 1B**). The eigenvectors of the solution represent the harmonic modes with the eigenvalues sorted in ascending order and reflecting the spatial frequency of the modes with lower modes representing lower spatial frequencies and higher modes representing higher spatial frequencies. Overall, the spatiotemporal activity can be perceived as a weighted contribution of these fundamental bases unfolding over the whole time recording for the spontaneous fMRI or as a weighted contribution of these fundamental bases reconstructing the task-based activations.

One of the features of functional connectivity is the surprisingly high functional connectivity between distant regions (14). We first investigated to what extent the different anatomical representations reconstruct the long-range connections. These were derived as an intersection of FC connections above 0.5 FC correlations and geodesic distance between the nodes above 40mm (**Figure 2A**). We then reconstructed these connectivity profiles with an increasing number of modes (1-200) derived from the four representative graphs (Geometry, EDR binary, EDR continuous and EDR+LR) (**Figure 2B**). The modes are ordered sequentially according to their spatial wavelength represented by their eigenvalues (i.e. mode 1 has the longest spatial wavelength). For all four graphs they monotonically decrease the reconstruction distance reaching on average 0.03 mse distance with about 20 modes and by 100 modes reach on average 0.01 mse distance before plateauing close to 0.005 mse distance on average for the full 200 modes. One noteworthy aspect is that much of the distance reconstruction happens between 0 and 20 harmonics suggesting that a small number of harmonics is responsible for most of the reconstruction. At 200 modes the EDR+LR outperforms the other spatial basis (Geometry, EDR continuous, EDR binary, paired t-test p<10^−4^). To assess the uniqueness of the LR connections within the EDR graph, we created a null model where we shuffled the LR connections in the EDR+LR graph representation. As expected, the specific rare long-range connectivity is important since the shuffled EDR+LR modes were unable to reconstruct the long-range functional connectivity to the same extent as the EDR+LR (**Figure S4**). Furthermore, to assess whether the EDR+LR performance is due to the unique combination of EDR and LR connectivity, we computed the reconstruction when using the structural connectome that implicitly contains the short-range and long-range connectivity, and long-range connectivity exceptions. However, as expected, the structural connectome graph representation showed less reconstruction capacity in comparison to the other representations (**Figure S5**). Lastly, to ensure robustness of the result, we carried out the analysis on an additional subset of 100 HCP participants reporting the same statistical significance between EDR+LR and Geometry (**Figure S11**).

**Figure 2.**
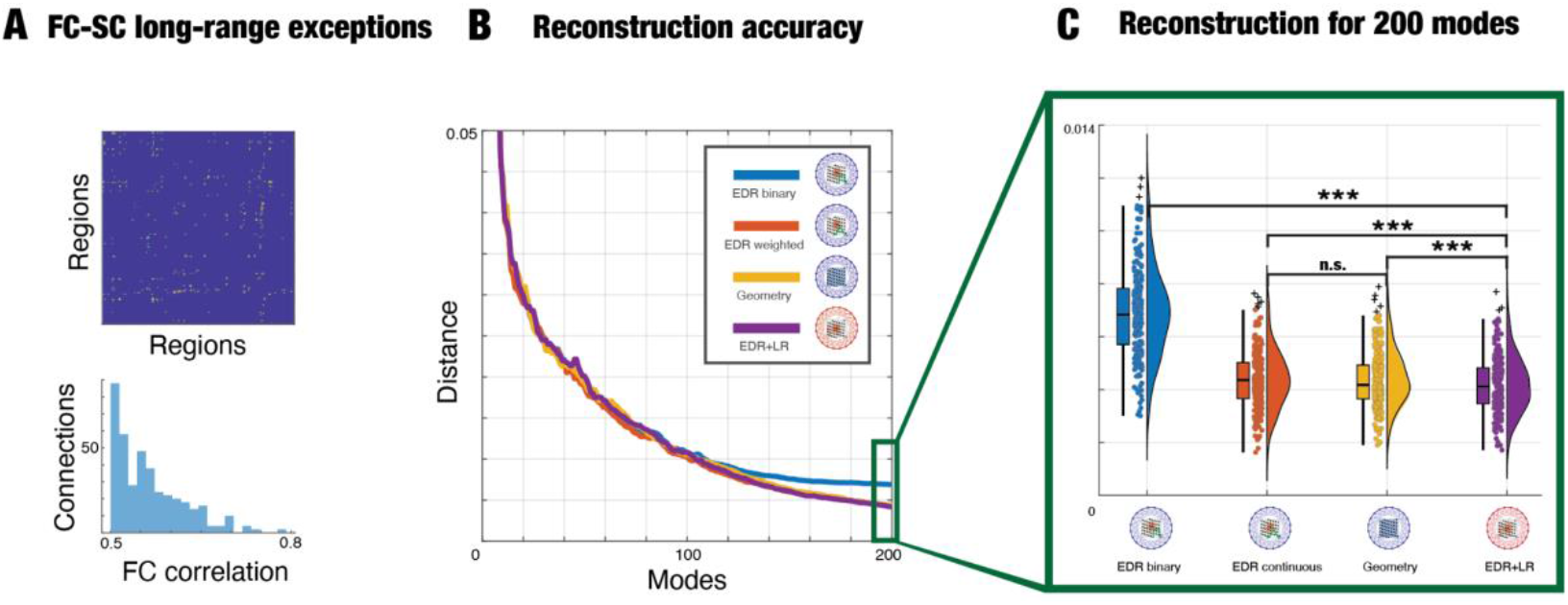
Better reconstructions of brain dynamics are found with EDR and rare long-range exceptions in the graph representation. **A)** One of the most important features of cortical dynamics are long-range functional connections (defined by high correlation values, >0.5 correlation, and Euclidean distance, >40mm). **B)** The reconstruction of FC long-range connections for an increasing number of modes (1-200) for the four representative graph representations. The individual lines show the average across all 255 HCP participants. **C)** EDR+LR is significantly better than the other graph representations when using a reconstruction with 200 modes as shown by the average result for the distance values across all the 255 HCP subjects (bonferroni corrected two-tailed paired t-test, EDR+LR and Geometry p<0.0005, EDR+LR and EDR continuous p<10^−4^, EDR+LR and EDR binary p<10^−4^, EDR continuous and Geometry n.s., * p<0.05, ** p<0.01, *** p<0.001).

### Less is more - EDR+LR reconstructs task with fewer modes

Using the same approach, we further investigated how well the different bases reconstruct the task-evoked brain activity from 255 healthy HCP participants. We used the 47 task-based contrasts derived from 7 HCP tasks each representing a different activation brain map and reconstructed them for an increasing number of modes (mode 1-200). For the 7 representative tasks the different bases demonstrate a similar monotonic pattern with steep fall in reconstructed mse distance before a slowdown with a near plateau-like behaviour around 200 modes and reconstructed mse distance values approximating 0.02 for most of the bases and tasks (**Figure 3A Top**). To analyse the reconstruction pattern, we computed the FC mse contribution of a given mode when added to the reconstruction. This demonstrates that the apparent bulk of the reconstruction is being obtained from a relatively small number of modes 0-20 in comparison to the rest (**Figure 3A Bottom**). This shows that reconstructing both spontaneous and task-evoked activity is represented in a very small space of 0-20 modes, suggesting that both types of dynamics, spontaneous and task-evoked, lie in a lower-dimensional manifold. Focusing only on the first 20 modes, we examined how the 47 task-evoked activations maps are reconstructed in comparison to the geometric modes. On average EDR+LR compared to Geometry shows the most accurate reconstruction across tasks up to 20 reconstructed modes (**Figure 3B**). By construction, the modes span an orthogonal basis set in which the individual mode contributions are mapped to. To motivate the neatness and accuracy of reconstructing the activation maps with as little EDR+LR as possible, we visually demonstrate the reconstruction of relational tasks for 5, 10, 15 and 20 modes showing the indistinguishable similarity to the activation map itself (**Figure 3C**). Moreover, it is not surprising that the EDR+LR basis, due to their unique topology, reconstructs with fewer modes more accurately the tasks as it can be appreciated in the motor tasks where more nuanced features are picked up in comparison to the geometric modes (**Figure 3D**).

**Figure 3.**
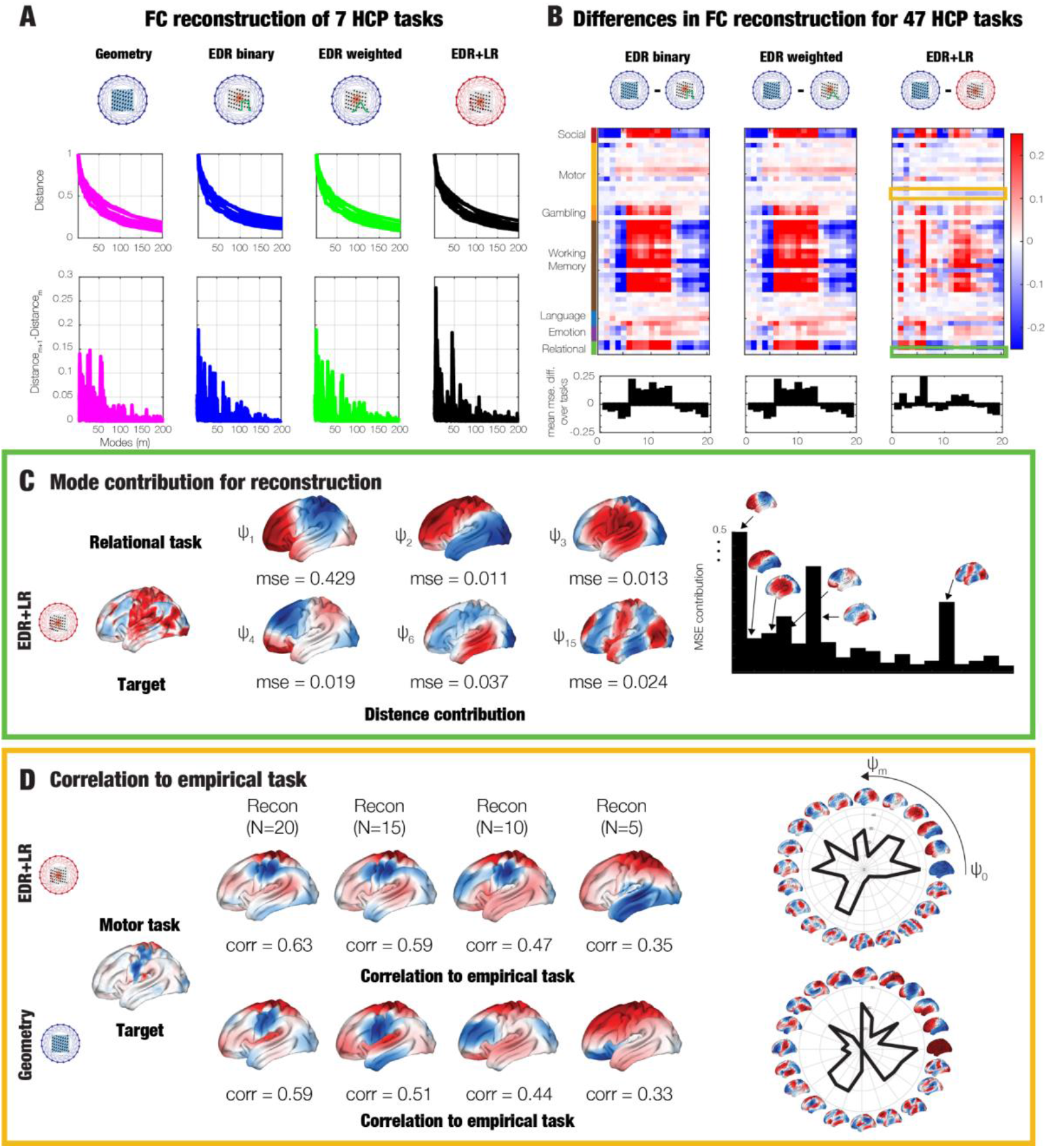
EDR+LR uses fewest harmonic modes to reconstruct task activity. **A)** For each of four graph representations (top panel) is shown the reconstruction of seven representative activation task fMRI maps in terms of normalised mse distance (distance normalised by the max of each task). As can be seen, lower frequency modes contribute disproportionately more toward the reconstruction distance as it can be seen by the elbow around 20 modes (lower panel). **B)** This can also be seen in the reconstruction mse distance for all 47 HCP tasks for the EDR+LR, EDR binary and EDR continuous, each benchmarked against the geometrical modes for the first 20 modes, where the top panel shows hues of blue with better performance of the EDR modes while red hues mean better performance of the geometric modes. The lower panel shows the average across the 47 HCP tasks. **C)** Individual mode contribution towards the reconstruction of the relational task. We show the disproportional contribution of some modes (1, 2, 3, 4, 6, 15) to the overall reconstruction, where the brain renderings show the mse distance reconstruction to the overall activation map (far left). **D)** Similarly, for the motor task target (far left), we compare the overall correlational contributions of the number of modes (using 20, 15, 10 and 5 modes) when using EDR+LR and geometry as the underlying representations. As can be seen, the reconstruction with EDR+LR converges more quickly for lower modes than geometry.

## Discussion

The unique mathematical formulation of harmonic modes links the description of how structure gives rise to the emerging spatiotemporal activity of brain dynamics. We show that EDR+LR modes have the smallest reconstruction distance for an increasing number of modes when describing the FC long-range connections of spontaneous fMRI activity. Furthermore, for the reconstruction of the 7 activation task fMRI maps lower frequency modes contribute disproportionately more toward the reconstruction error. We therefore reconstructed the error for the 47 HCP tasks benchmarked against the geometrical modes for the first 20 modes. On average EDR+LR showed the most accurate reconstruction across tasks and number of reconstructed modes 1-20. Our results demonstrate the importance of long-range connectivity as a key feature of shaping brain functional activity both for the spontaneous and task-based fMRI. Moreover, functional brain activity is shown to be on a lower-dimensional manifold span by a subset of these fundamental modes with the most appropriate representation from the EDR+LR graph, suggesting that less is more for efficient information processing in the brain.

In both spontaneous and task-based reconstruction cases, the EDR+LR demonstrate high reconstruction only with a subset of modes from its harmonic repertoire. Despite the overall better performance of the EDR+LR harmonic modes, it is remarkable that the other harmonic bases, geometric and EDR-based, performed strongly as well. This excellence can be seen from the fact that all four reconstruction schemes are able to predict behavioural measures fluid intelligence and participant’s processing speed. The results show that this prediction is driven by the brain state (task-evoked over spontaneous fMRI) consistently across the four graph representations (**Figure S6-8**). This reflects a fundamental insight where large-scale brain organisation can be described as lying in a low-dimensional manifold. This in part can be explained by the brain’s coordinated cognition and behaviour which cannot happen without integrative tendencies of its underlying dynamics. Indeed, brain dynamics operating in a reduced number of dimensions have been shown to predict more effectively the brain’s behaviour (15). As such one can talk of brain activity as a flow on this low dimensional manifold embedded in the space of these relatively few harmonic modes (16).

One of the fundamental considerations is what type of brain’s dynamics we wish to reconstruct. Unlike the traditional approach where the whole static functional connectivity is reconstructed (5), we focused on reconstructing the most salient features of the brain’s spontaneous fMRI activity, namely the functionally strong long-range connection. Our work underscores the cardinal role of long-range connectivity in cognitive processing and advocates for prioritising the reconstruction of exceptional connections over exhaustive coverage of the entire functional connectivity matrix. With similar logic, we did not regress out the global signal from the spontaneous fMRI as we consider the global and fluctuating fMRI activity an important feature of emergent network effects of interacting non-linear regional dynamics (17). As expected, when we computed the analysis, applying global signal regression, the reconstruction of EDR+LR and Geometry were statistically non-significant (**Figure S9**). Moving beyond, it is important to consider temporally evolving descriptions of brain dynamics as recent work has demonstrated the relevance of dynamics in understanding brain function and its related pathologies (18). Also, many whole-brain modelling techniques have been suggesting the need to consider further descriptors of brain activity that goes beyond the static FC description (19). Ultimately, as recently suggested by the spatiotemporal neuroscience of Northoff and colleagues (20, 21), the brain’s dynamic spatiotemporal organisation might reveal the link between the neuronal and mental features to elucidate concepts such as consciousness, self and time speed perception.

Flexible human cognition and behaviour reflect a highly dynamic balance of functional integration and segregation. This in turn is supported by the rich topology of the structural connectome (22). A growing body of literature has shown that these dynamics are poised at the edge of criticality, a dynamic regime with long-range spatial and temporal correlations in which information can be optimally processed (19). This is consistent with a novel computational framework by Jaeger and colleagues (23), suggesting that an understanding of computing comes from an understanding of the structuring of processes, rather than how classical models of computing systems describe the processing of structures. They also stress how this can come via an understanding of modelling physical computing systems bottom-up, which is the main aim of the investigation here, where the topology of the computing system, here the brain, shapes the near-critical dynamics of the system. In the brain, the rare long-range structural connections are some of the key anatomical features supporting time-critical information processing. Their spatially specific location has been linked to the emergence of known resting-state networks and are important for task-based processing (12). We therefore hypothesise that evolutionary pressures are likely to have refined EDR connectivity with long-range exceptions enabling more complex cognitive functions. This hypothesis should be investigated in future cross-species studies.

In this work, we derived both EDR binary and EDR continuous harmonic modes. These reflect different methodological considerations when calculating the Laplacian eigenmaps (7). We have applied the continuous form of the graph Laplacian on the EDR (EDR continuous) showing that this simple change improves the reconstruction accuracy by about 0.0025 distance to the binarized version (EDR binary) making it practically on the same footing as the geometric bases (EDR continuous and geometry are not statistically different from each other). Reassuringly, recent work has reported similar observations when comparing the non-binarised structural connectomes graph representation to the geometric modes (24). It is therefore warranted to unify the methodological approaches before comparing the superiority of the different anatomical features as the differences might be simply explained by methodological choices themselves. Therefore, we caution future research to unify the applied methodologies in this direction.

Given the relevance of rare long-range connectivity for complex brain dynamics in humans, it is also important to consider how these findings might translate to other non-human species. Unfortunately, a direct comparison between species is challenging due to the different methodologies used. Unlike non-human species studies that use track tracing studies to describe the anatomical connectivity of the brain, human experiments rely on non-invasive techniques like diffusion MRI. Furthermore, the challenges in estimating long-range connectivity via dMRI further complicates the direct comparison (25, 26). Yet, a growing body of non-human species studies have converged on a general principle that smaller brains such as the mice brain are denser, all-to-all connected, whereas larger brains such as the primate brains are sparser with weak long-range connections reflecting further regional specialisation (27, 28). Therefore, it can be hypothesised that long-range EDR connections together with the rare connectivity exceptions play an important role in the emergence of complex computational capabilities. This opens up exciting future cross-species research of the impact of rare long-range connections on the brain’s computational capabilities.

Understanding rare long-range connections’ impact on the emergent brain dynamics will also help clinical diagnosis in neuropsychiatry and neurology and inform novel clinical treatments. For instance, the weak nature of these rare long-range connections might be abnormally affected in disconnection syndromes such as Alzheimer’s disease and schizophrenia and this in turn might have a disproportionate impact on the large-scale emergent dynamics affecting cognition and behaviour (27). Moreover, novel treatment solutions, such as transcranial electrical stimulation, will rely on model optimization where anatomical connectivity plays an important role (29, 30). In the future, the specific inclusion of rare long-range connections in the models might ensure more accurate description of the disorders as well as more efficient stimulation protocol for possible treatments.

## Materials and Methods

### Experimental Data

#### HCP Functional MRI

We used the publicly available Human Connectome Project (HCP) dataset, Principal Investigators: David Van Essen and Kamil Ugurbil: 1U54MH091657) with the funding coming from sixteen NIH Institutes and Centres supporting the NIH Blueprint for Neuroscience Research; and by the McDonell Centre for Systems Neuroscience at Washington University. All participants joined voluntarily and provided informed consent. The open-access data used in this study were obtained through the WU–Minn HCP consortium, following approval from the local ethics committee. The data were shared with the authors in accordance with the terms specified by the HCP for data usage. All procedures conducted in this study adhered to the protocols outlined in these data use terms. For a comprehensive description of the image acquisition protocol, preprocessing pipelines (31), and ethics oversight, please refer to the detailed account provided (31, 32).

### Spontaneous fMRI dataset

We used the spontaneous fMRI dataset from the freely accessible database with connectome DB account at https://db.humanconnectome.org. Timeseries were minimally processed. Consistent with work of Pang and colleagues (5), we used a subset of 255 participants (22-35yo, 132 F and 123 M) who completed all spontaneous and tasks-based fMRI recordings, further excluding twins and siblings. For the auxiliary dataset we used a subset of 100 HCP participants which were different to the main analysis performed with the 255 HCP participants. The neuroimaging acquisition was carried out on a 3-T connectome-Skyra scanner (Siemens). A single spontaneous fMRI acquisition, lasting approximately 15 minutes, was conducted on the same day. During this session, participants kept their eyes open with relaxed fixation on a projected bright crosshair against a dark background. The HCP website offers comprehensive details on participant information, acquisition protocols, and data preprocessing for both spontaneous and the seven tasks. In summary, the data underwent preprocessing using the HCP pipeline, which employs standardised methods with FSL (FMRIB Software Library), FreeSurfer, and Connectome Workbench software. This standardised preprocessing encompassed correction for spatial and gradient distortions, head motion correction, intensity normalisation, bias field removal, registration to the T1-weighted structural image, transformation to the 2-mm MNI space, and application of the FIX artefact removal procedure. Head motion parameters were regressed out, and structured artefacts were removed using independent component analysis, followed by FMRIB’s ICA-based X-noiseifier (ICA+FIX) processing. The preprocessed time series for all grayordinates were in the HCP CIFTI grayordinates standard space, available in the surface-based CIFTI file for each participant during spontaneous fMRI. Lastly, for Figure S9, we also regressed out the global signal before carrying on with further analysis on the spontaneous fMRI.

### Tasks-based fMRI dataset

For the task-based fMRI analysis, we obtained fMRI data from 7 distinct task domains known to reliably engage a diverse range of neural systems (5, 31). The tasks included were social, motor, gambling, working memory (WM), language, emotion, and relational. We used the specific contrasts within each task domain, highlighting the key contrast investigated in this study. These contrasts were provided by work of Pang and colleagues (5) from https://osf.io/xczmp/ in “S255_tfMRI_ALLTASKS_raw_lh” .mat file. In total, the analysis encompassed 47 contrasts, incorporating the 7 key contrasts. In brief, the analysis was performed on individual task-activation maps generated through FSL’s cross-run (Level 2) FEAT analysis (33). The task maps, provided by the Human Connectome Project (HCP), were used with minimal smoothing (2 mm), and mapped onto the fsLR-32k CIFTI space. This mapping was achieved using multimodal surface matching, resulting in a representation of each individual’s task data (32,492 vertices). Additional information about each task and contrast as well as further details on the data are provided elsewhere (5, 31). The task-evoked fMRI reconstruction distance was computed on the parcellated activation maps, unlike those using spontaneous fMRI, where the reconstruction distance was performed on the parcellated functional long-range connections.

### fMRI parcellation

A custom MATLAB script, utilising the ‘ft_read_cifti’ function from the Fieldtrip toolbox, was employed to extract the average time series of all grayordinates in each region defined by the Glasser360 parcellations (180 regions per hemisphere) in the HCP CIFTI grayordinates standard space. For each hemisphere the vertex-space to ROI-space meant going from 32,492×1200 to 180×1200 for spontaneous fMRI and 32,492×1 to 180×1 for task-based fMRI. Consistent with work by Pang and colleagues (5) our analysis focused on the left hemisphere only.

### HCP Diffusion MRI

To obtain the structural connectivity for the fitting of the EDR and derivation of long-range exceptions to the EDR, we used the high-resolution connectivity maps from dMRI tractography (34). These were provided by work of Pang and colleagues (5) in “S255_high-resolution_group_average _connectome_cortex_nomedial-lh” .mat file. In brief, the connectome was derived by estimating the connectivity of each of the 32,492 vertices within the cortical surface mesh by tracing streamlines from each point until they terminated at another point. Connection weights between vertices, treated as nodes, were determined as the number of interconnecting streamlines without normalisation (35). The dMRI tractography was conducted on individuals from the Human Connectome Project (HCP). Subsequently, the individual weighted connectivity matrices were combined, each of size 32,492 × 32,492, to generate a group-averaged connectome. The weights in this connectome represented the average number of streamlines, providing a comprehensive depiction of group-level connectivity. Further details can be found in previous publication by Pang and colleagues (5).

### Structural MRI

For the fitting of the EDR, we used the Euclidean distance (on a flattened cortical surface) between the vertices of the cortical mesh representation for the left hemisphere (32,492×32,492). This mesh was derived from the FreeSurfer’s fsaverage population-averaged template available on github.com/ThomasYeoLab/CBIG/tree/master/data/templates/surface/fs_LR_32k. It is to be noted, we used the version provided by Pang and colleagues (5) in the “fsLR_32k_midthickness-lh” .vtk file.

### Exponential Distance Rule (EDR)

Previous work has demonstrated that the brain white-matter wiring, based on retrograde tract tracing in non-human primates, can be analytically approximated by the Exponential Distance Rule (EDR) (6). Here, we derived the Exponential Distance Rule of the underlying human anatomy using diffusion MRI (**Figure S3**). Mathematically, the exponential distance rule can be described with exponential decay function as follows:

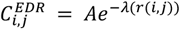

where *r*(*i, j*) is the geodesic distance between vertices *i* and *j* and *λ* is the decay. Consistent with previous literature, we estimated the parameters (*A* and *λ*) for the exponential decay model using a least-squares method as follows *y* = *Ae*^−*λx*^, where *y* represents a mean connection weight of a given geodesic distance and *x* represents the given geodesic distance (12). In detail, we have generated 400 bins of equal geodesic distance taking the bins spanning 10mm to 170mm (thus excluding the first 25 bins in the fitting procedure). The estimation yielded *A* = 0.066 and *λ* = 0.162 mm^−1^ where the exponential decay parameter lambda is consistent with previous literature (10, 12). We used this estimation for the construction of the EDR+LR graph. For the EDR binary and EDR continuous, we used previously reported exponential decay parameter of *λ* = 0.12 mm^−1^ to be consistent with work of Pang and colleagues (5) (See sections EDR binary and EDR continuous).

### Relationship to Belkin and Niyogi

The exponential distance rule, as an optimal solution for connecting distance-separated brain regions in the brain, can also be intuitively understood from first principles. Belkin and colleagues have analytically shown the relationship between graph Laplacian, Laplace Beltrami Operator and the heat kernel which is the optimal solution for locality preservation - formally as 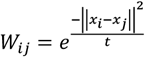 where *t* is the decay parameter of the heat kernel (7). It can thus be appreciated that this equation also follows exponential decay (gaussian) similar to the EDR.

### Harmonic Modes

In this work, we used four different types of graph representations to describe various aspects of anatomical features or methodological approaches. Namely, we carried out the analysis on what we call Geometric, EDR binary, EDR continuous and EDR+LR modes. In what follows, we describe the remaining three types of harmonic modes representations.

#### EDR binary

For the EDR binary, we use the EDR with the same parameters as in work of Pang and colleagues (5) to define the weight of a given edge between vertices *i* and *j*. In other words, the weight is determined by the geodesic distance between regions *i* and *j* and the fitted lambda parameter, *λ* = 0.12 mm^−1^ (see section Exponential Distance Rule). Then, as in work by Pang and colleagues (5), we created a binary adjacency matrix where nodes *i* and *j* are retained and binarized only if the weight strength surpasses randomly distributed distribution of the weights. This option results in a binary adjacency matrix whereby *C*_*ij*_ = 1 if i and j are above randomly distributed distribution of the weights and *C*_*ij*_ = 0 if i and j are below the randomly distributed distribution of the weights. The choice of this approach was motivated to stay consistent with previous work by Pang and colleagues (5) in order for the results to be directly comparable.

#### EDR continuous

For the EDR continuous, we similarly use the EDR with the same parameters to define the weight of a given edge between vertices *i* and *j* using the EDR with *λ* = 0.12 mm^−1^. Unlike the thresholding in EDR binary (applied in work of Pang and colleagues (5)) where connections are retained and binarized if they surpass connection weights from a randomly derived distribution, here all the connections and their weights are kept. This option results in a weighted adjacency matrix whereby *W*_*ij*_ = *Ae*^−*λ*(*r*(*i,j*))^. Furthermore, we argue in this paper that this detailed explanation between EDR binary and EDR continuous adjacency matrices is warranted as it zeroes in on what is an appropriate comparison between graph Laplacian, and continuous Laplace-Beltrami analysis and we motivate future comparative research in this direction.

#### Geometry

The geometric modes were calculated using the Laplace Beltrami Operator (LBO) on the cortical mesh. We used the publicly available version from previously published work by Pang and colleagues which can be downloaded from https://osf.io/xczmp/ in “fsLR_32k_midthickness-lh_emode_200” .txt file (5). In brief, the LBO is in general defined as follows:

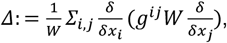

With *g*^*ij*^ being the inverse of the inner product metric tensor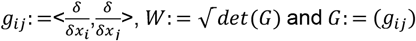. The solution of the eigenvalue problem was implemented in a python package LaPy using the cubic finite element method (36). For further details consult work by Pang and colleagues (5). Although not explicitly stated, the derivation leverages an exponential kernel that is reminiscent of the EDR.

#### EDR+LR

Previous research has shown that human as well as non-primate anatomy is characterised by a relatively small proportion of long-range outliers to the EDR (10, 12). Therefore, for the EDR continuous adjacency matrix we wanted to implement a version where these long-range (LR) exceptions are taken into account. Using the structural connectivity matrix, we computed the binned distribution (400 bins) as a function of geodesic distance. We defined connectivity exceptions as 3 standard deviations above the mean for a given distance bin that are longer than 40mm (**Figure S2**). To derive the EDR+LR connectivity matrix we combine the EDR continuous with LR exceptions to the EDR. Moreover, we also created a shuffled EDR+LR where the locations of the LR were randomly assigned in the connectivity matrix (**Figure S4)**.

### EDR+LR relationship to Connectome Harmonics

Combining short-range and long-range connectivity can be performed in many ways. Indeed our previous work on connectome harmonics has defined the anatomical connectivity in terms of short-range, nearest-neighbour connections on the cortical surface, combined together with long-range connections, derived from the diffusion MRI in terms of the connectome (4). In this light, here, we derive the short-range connections in a more principled way through the “EDR continuous” while accounting for the long-range connections in terms of the exceptions to the EDR as stated above. Furthermore, we avoid binarization of the adjacency matrix for the calculation of the Laplacian as it has shown to retain important information in the reconstruction of both spontaneous and task-evoked fMRI from our results on binary and continuous EDR brain modes.

### Laplacian Decomposition

Having derived the EDR+LR, EDR binary and EDR continuous adjacency matrix, we calculated the normalised graph Laplacian as

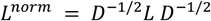

with *L* = *D* − *A* where D is the diagonal degree matrix defined as 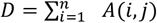 Finally, the harmonic modes were computed as eigenvectors of the following eigenvalue problem

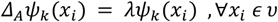

with *λ*_*k*_, *k ϵ* 1, …, *n* are the eigenvalues of *Δ*_*A*_ and *ψ*_*k*_ is the *k*^*th*^ harmonic mode. We report visually the harmonic modes for EDR+LR, Geometry, EDR continuous, EDR binary rendered on the brain (**Figure S1**).

### Decomposition of brain activity with harmonic modes

We can represent the spatiotemporal spontaneous fMRI recording and the activation maps of task-based fMRI as a weighted contribution of the harmonic modes as follows

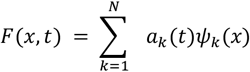

Where *F* is the spatiotemporal timercordings for each subjects with dimension 32,492×1200 (*x, t*), *a*_*k*_(*t*) has dimension 1×1200 and is the contribution of *k*^*th*^ harmonic to the F timecourse at time t. Note that for the purely spatial data of task-based fMRI the same applies except of the contributions being independent of time ie *a*_*k*_(*t*) → *a*_*k*_. Both in spontaneous and task-based fMRI, the contributions are computed as the inner product between the spatial patterns and harmonic modes

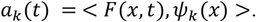

### Reconstruction error

To compare both the spontaneous and task-based empirical fMRI data with the reconstructed data with a subset of harmonic modes, we first parcellated the data to Glasser360 parcellation (we focused on the left hemisphere resulting in 180 nodes). For the spontaneous fMRI, we calculated the interregional functional connectivity (FC −180×180) and focused on the most salient features by reconstructing the long-range functional connectivity derived as a subset of connections with high-correlation values (> 0.5 correlation) and a long Euclidean distance (> 40mm). Then, we calculated the reconstruction error as the mean squared error (mse) distance between the empirical and reconstructed long-range functional connectivity. For the task-based fMRI we calculated the reconstruction error as the mse distance between the empirical and reconstructed activation maps. Lastly, for the behavioural analysis of **Figure S6-8**, we correlated the reconstruction (for 200 modes) of spontaneous and task-evoked activity with fluid intelligence of participants in terms of three variables in the HCP data, namely 1) the number of correct responses in the PMAT24 A test and 2) processing speed in terms of Pattern Completion Processing Speed (CardSort_UnAdj and ProcSpeed_Unadj) (37).

## Acknowledgments

Jakub Vohryzek is supported by EU H2020 FET Proactive project Neurotwin grant agreement no. 101017716, Yonatan Sanz-Perl is supported by ‘ERDF A way of making Europe’, ERDF, EU, Project NEurological MEchanismS of Injury, and Sleep-like cellular dynamics (NEMESIS; ref. 101071900) funded by the EU ERC Synergy Horizon Europe, Morten L. Kringelbach is supported by the European Research Council Consolidator Grant: CAREGIVING (615539), Pettit Foundation, Carlsberg Foundation and Center for Music in the Brain, funded by the Danish National Research Foundation (DNRF117). Gustavo Deco is supported by the Spanish Research Project PSI2016-75688-P (Agencia Estatal de Investigación/Fondo Europeo de Desarrollo Regional, European Union); by the European Union’s Horizon 2020 Research and Innovation Programme under Grant Agreements 720270 (Human Brain Project [HBP] SGA1) and 785907 (HBP SGA2); and by the Catalan Agency for Management of University and Research Grants Programme 2017 SGR 1545.

