## Supplementary Information for "Beyond cortical geometry: brain dynamics shaped by rare long-range connections"

### **Supporting Information for:** Beyond cortical geometry: brain dynamics shaped by rare long-range connections

Jakub Vohryzek[1,2], Yonatan Sanz-Perl[1], Morten L. Kringelbach[2,3,4] and Gustavo Deco[1,5]
1. Centre for Brain and Cognition, Computational Neuroscience Group, Department of Information and Communication Technologies, Universitat Pompeu Fabra, Barcelona, Spain
2. Centre for Eudaimonia and Human Flourishing, Linacre College, University of Oxford, UK
3. Department of Psychiatry, University of Oxford, Oxford, United Kingdom
4. Centre for Music in the Brain, Aarhus University, Aarhus, Denmark
5. Institució Catalana de la Recerca i Estudis Avançats (ICREA), Barcelona, Spain
* Jakub Vohryzek

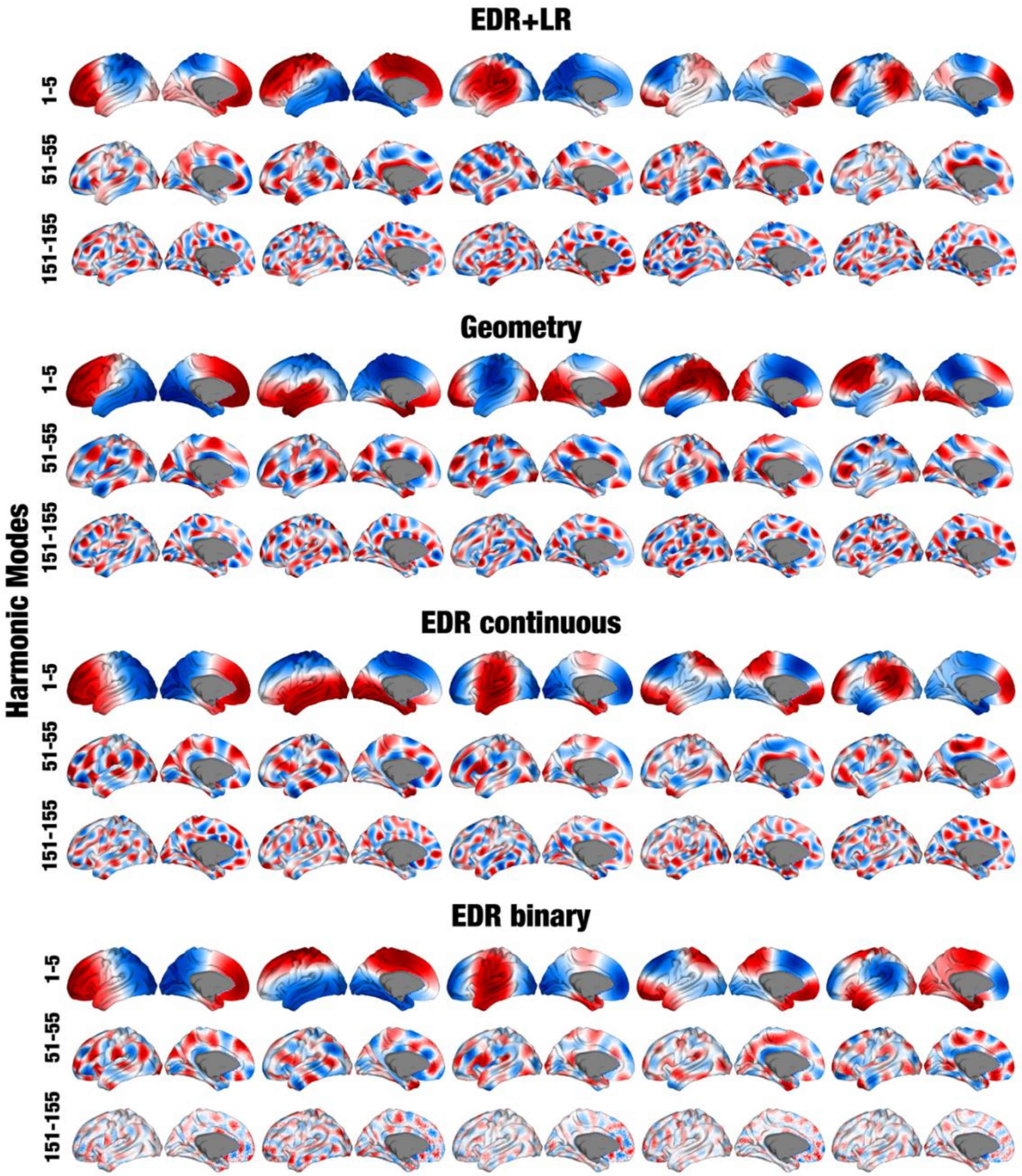


**Fig. S1. Brain renders of harmonic modes for EDR+LR, Geometry, EDR continuous, EDR binary.** Visual comparison of the harmonic modes for different graph representations for varying degrees of spatial frequency. Here, we chose to plot ranges of 5 harmonic modes (1-5, 51-55, 151-151) moving from global representation to more localised representation of brain activity while still reflecting the underlying graph representation. Already, at the most global range (1-5 harmonic modes), a fundamental difference can be observed between the different graph representations. At the highest spatial frequency (151-155) the impact of binarization to the EDR binary graph representation can be observed reflecting the underperformance compared to the other graphs.

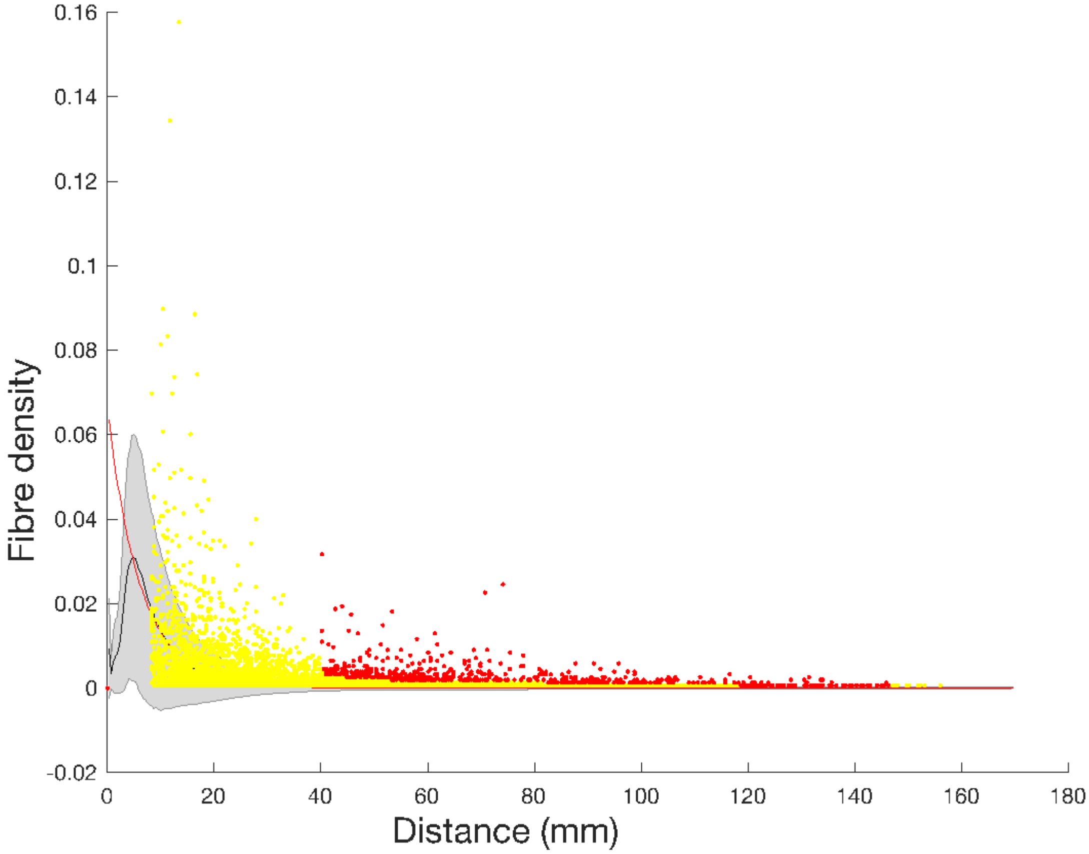


**Fig. S2. Long-range exceptions derived from the Exponential Distance Rule (EDR).** A histogram showing the long-range exceptions to the exponential distance rule in red. These long-range exceptions are derived by considering EDR fiber density connections stronger than three standard deviations for a given distance bin and being longer than 40mm. An exponential curve (in red) was fitted to the connectivity distribution of the structural connectome as a function of geodesic distance with the mean represented in black and standard deviation in shaded grey. The yellow dots represent the individual connections overlay on the connectivity distribution above 10mm.

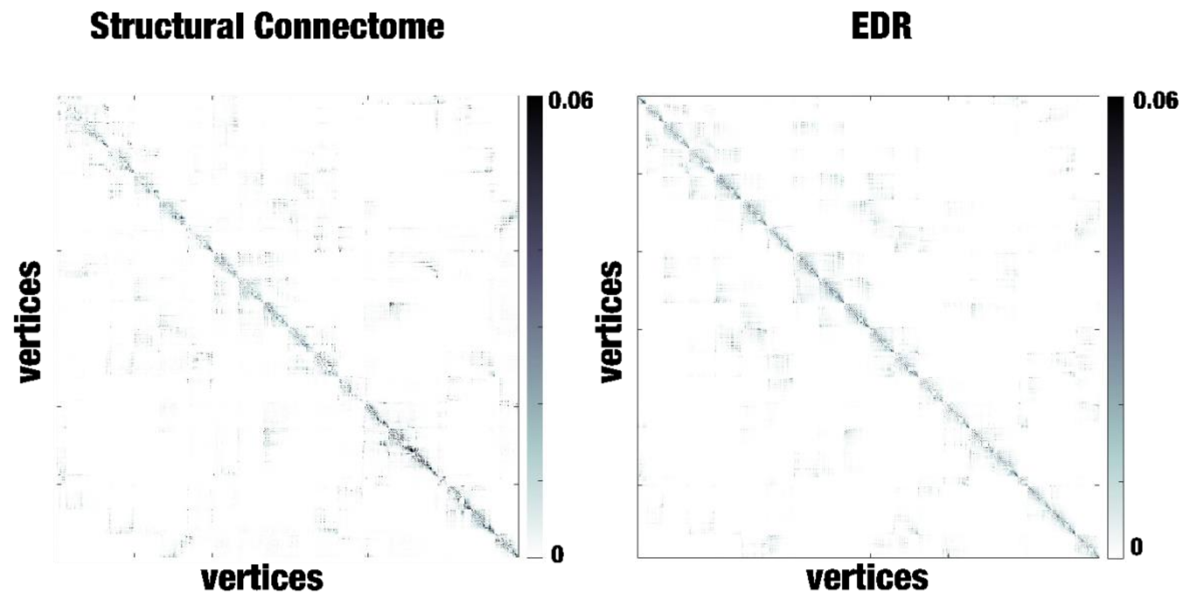


**Fig. S3. Comparison between the Structural connectome and its fitted exponential distance rule (EDR).** A visual representation of the structural connectome and the EDR connectome appreciate the similarity between the two. The key differences are the rare long-range connections derived as fiber density connections stronger than three standard deviations for a given distance bin and being longer than 40mm.

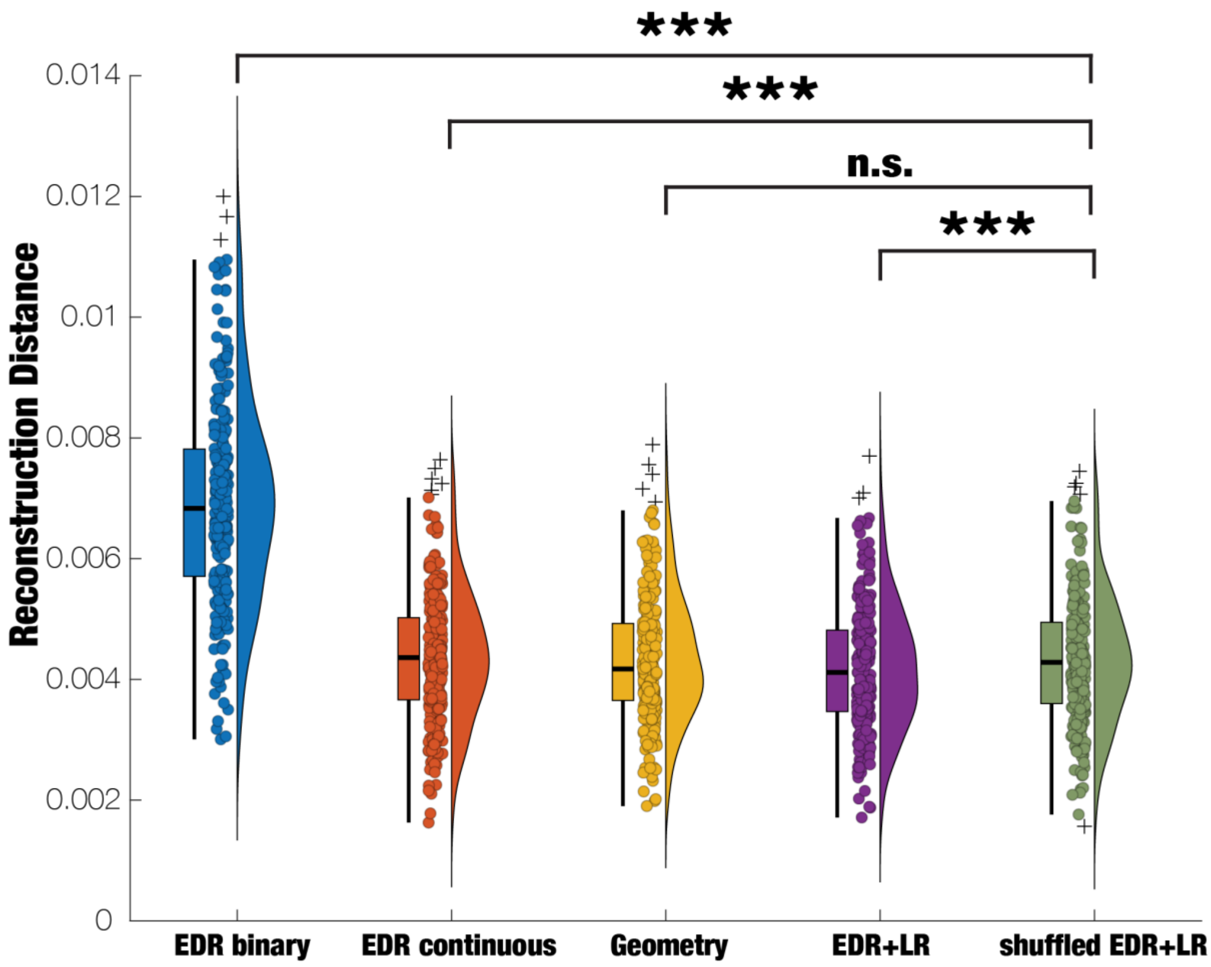


**Fig. S4. As expected, shuffling EDR+LR modes were as good as Geometry but significantly less good at reconstructing the long-range functional connectivity as the EDR+LR.** The reconstruction of FC long-range connections (defined by high correlation values, >0.5 correlation, and Euclidean distance, >40mm) for 200 modes using all four graph representations plus null model with shuffled LR connections of the EDR+LR graph representation (shuffled EDR+LR). Shuffled EDR+LR has significantly higher MSE reconstruction compared to the EDR+LR, and lower MSE for other EDR graph representations (SC vs. EDR+LR $p<10^{-3}$, SC vs. EDR binary $p<10^{-3}$, SC vs. EDR continuous $p<10^{-3}$, SC vs. Geometry p = n.s., Bonferroni corrected two-tailed paired t-test, Bonferroni corrected two-tailed paired t-test, * $p<0.05$, ** $p<0.01$, *** $p<0.001$).

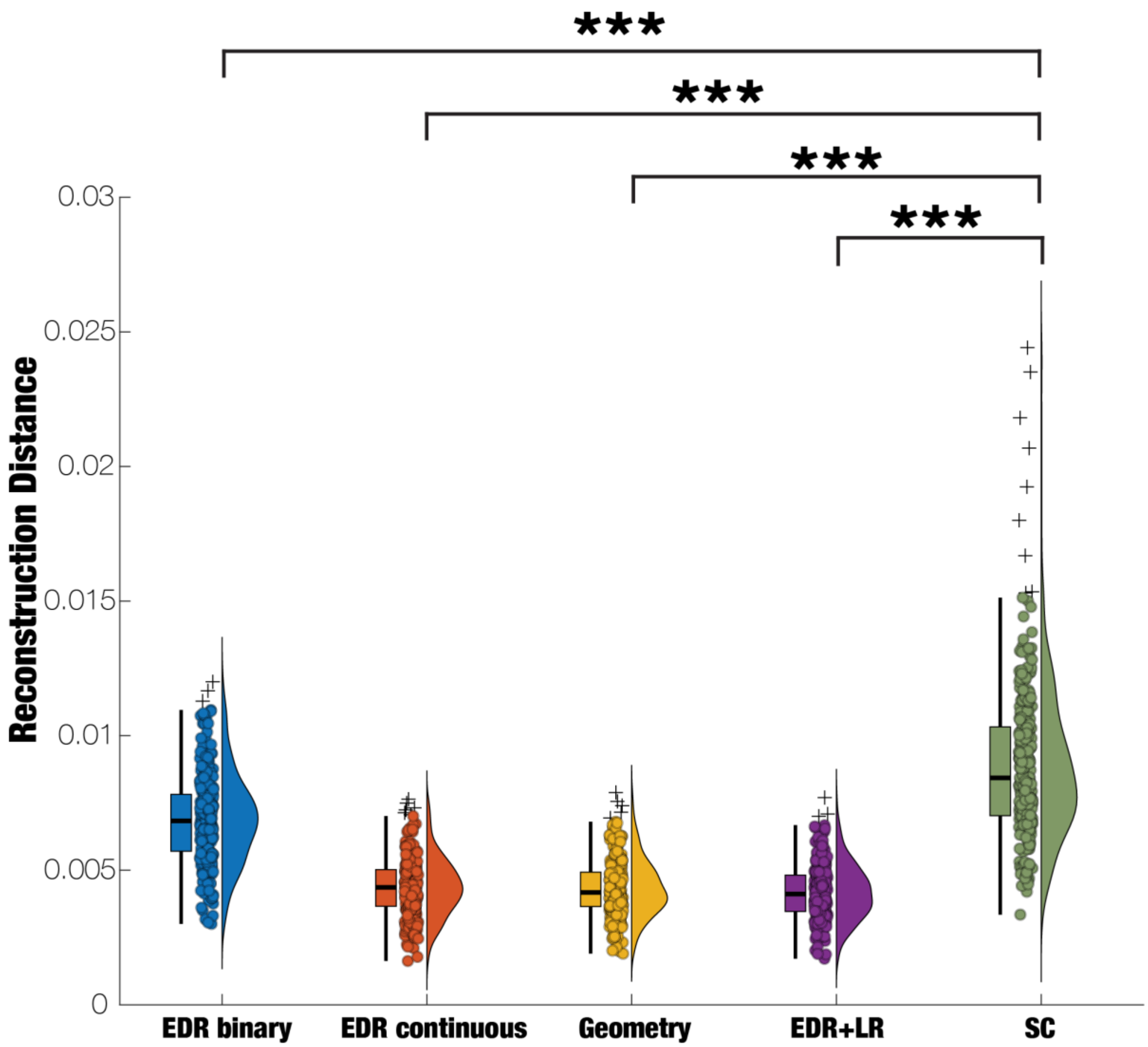


**Fig. S5. Reconstruction with structural connectome reconstructs significantly less of long-range functional connectivity of spontaneous brain activity compared to the other anatomical constraints.** The reconstruction of FC long-range connections (defined by high correlation values, >0.5 correlation, and Euclidean distance, >40mm) for 200 modes using all four graph representations plus state-of-the-art dense structural connectome (SC). SC has significantly higher MSE reconstruction compared to the other graph representations (SC vs. EDR binary $p<10^{-3}$, SC vs. EDR continuous $p<10^{-3}$, SC vs. Geometry $p<10^{-3}$, SC vs. EDR+LR $p<10^{-3}$, Bonferroni corrected two-tailed paired t-test, Bonferroni corrected two-tailed paired t-test, * $p<0.05$, ** $p<0.01$, *** $p<0.001$).

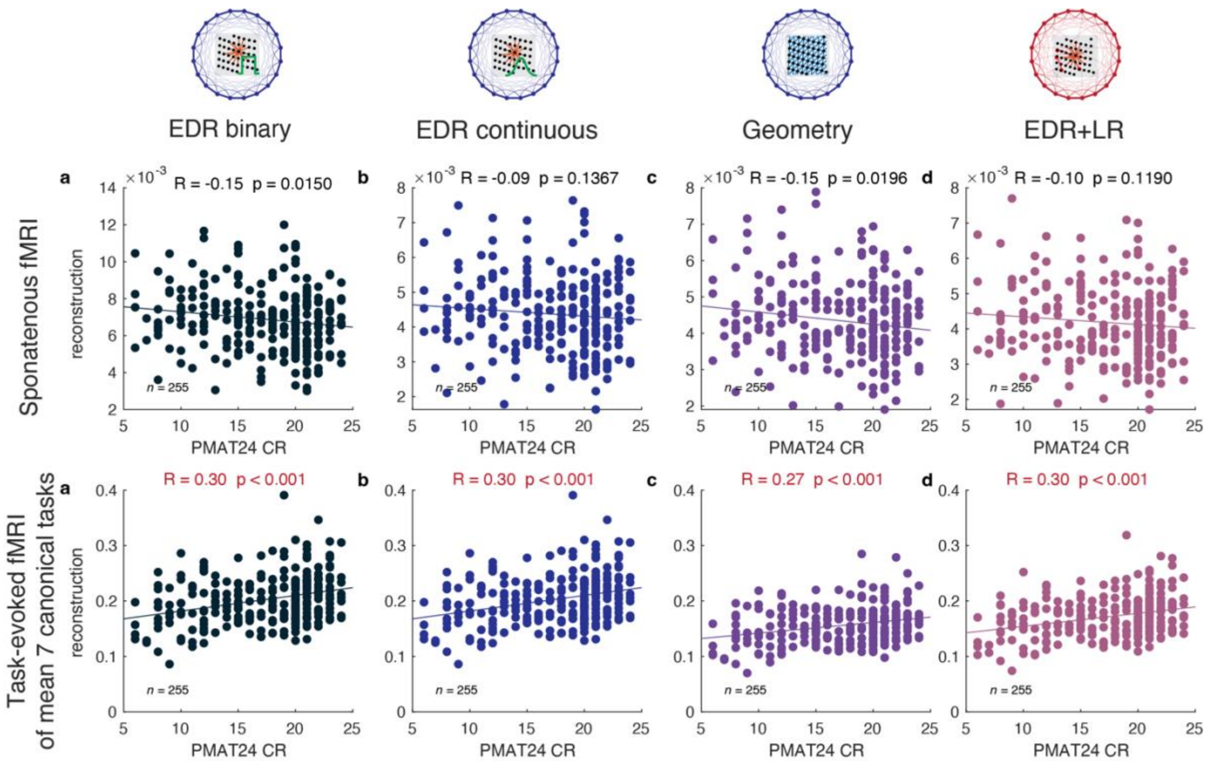


**Fig. S6. All reconstruction schemes correlate with a measure of fluid intelligence (PMAT24 CR) in task fMRI but not for spontaneous fMRI**. *Row with spontaneous fMRI: a-d* - MSE reconstruction and PMAT24 CR for EDR binary, EDR continuous, Geometry and EDR+LR respectively. *Row with task-evoked fMRI of mean 7 canonical tasks: a-d* - MSE reconstruction and PMAT24 CR for EDR binary, EDR continuous, Geometry and EDR+LR respectively.

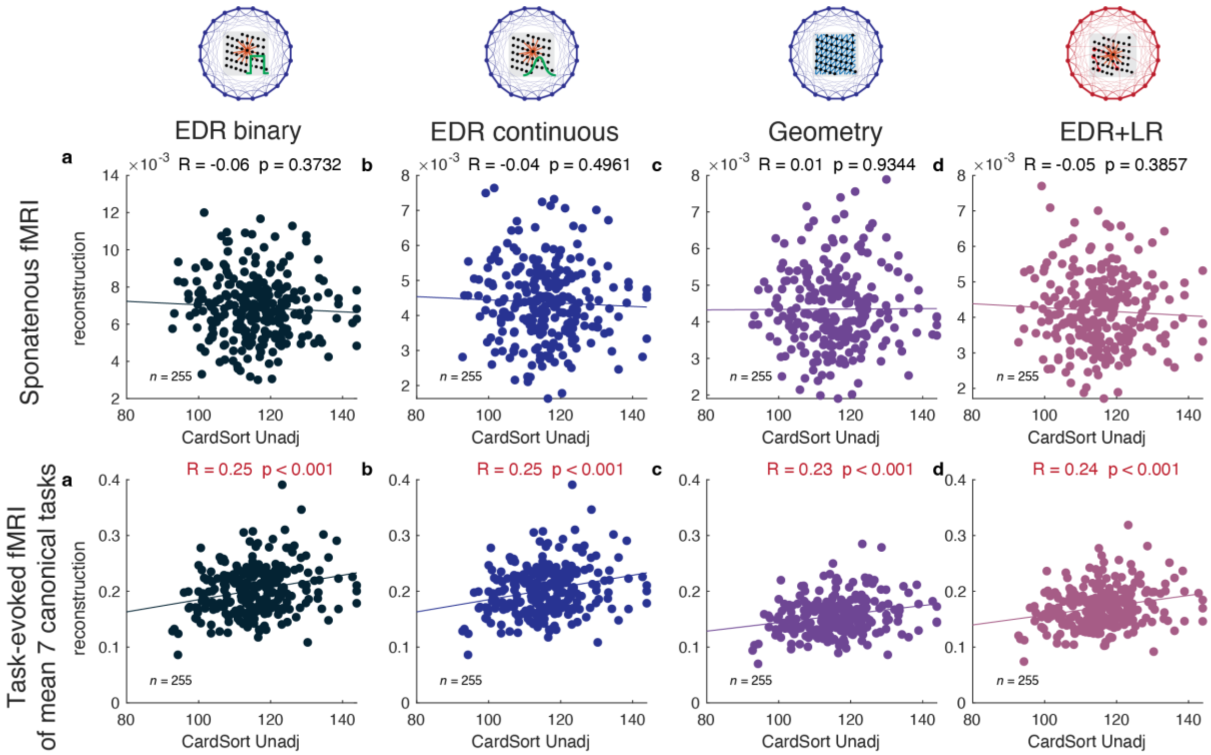


**Fig. S7. All reconstruction schemes correlate with a measure of Processing Speed (CardSort Unadj) in task fMRI but not for spontaneous fMRI.** *Row with spontaneous fMRI: a-d-* MSE reconstruction and PMAT24 CR for EDR binary, EDR continuous, Geometry and EDR+LR respectively. *Row with task-evoked fMRI of mean 7 canonical tasks: a-d* - MSE reconstruction and PMAT24 CR for EDR binary, EDR continuous, Geometry and EDR+LR respectively.

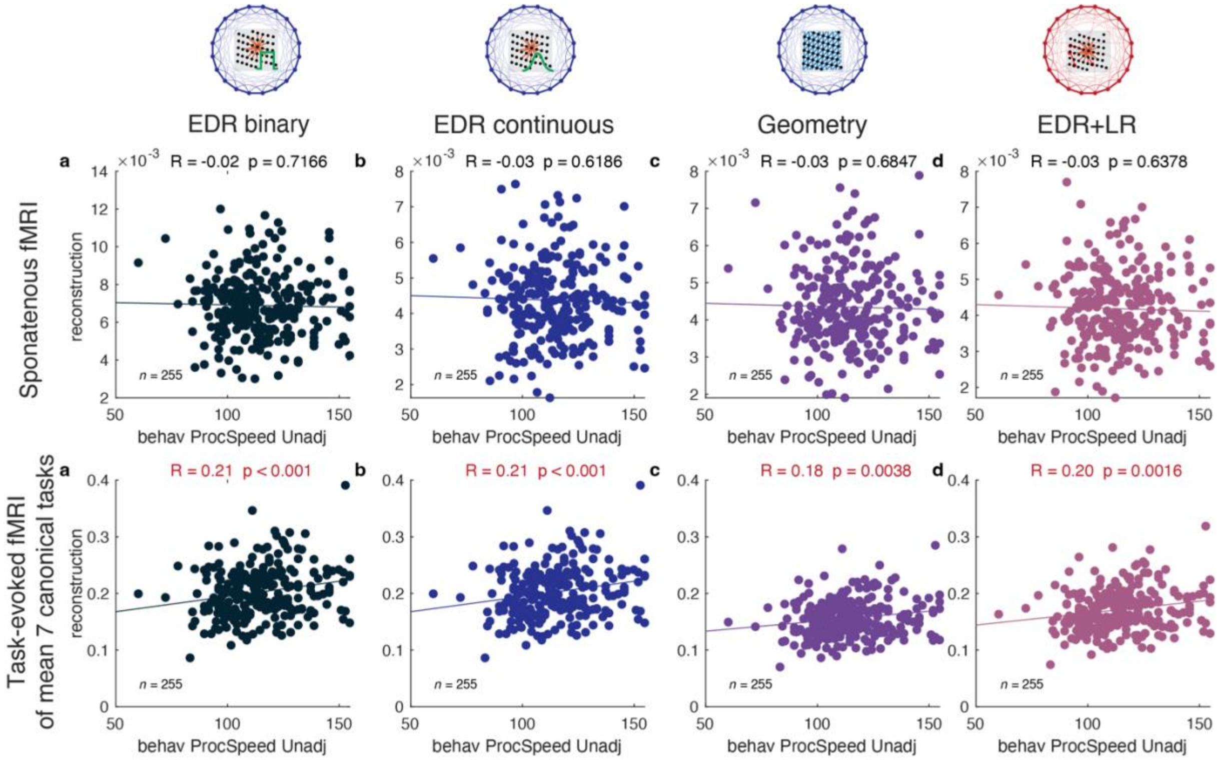


**Fig. S8. All reconstruction schemes correlate with a measure of Processing Speed (ProcSpeed Unadj) in task fMRI but not for spontaneous fMRI.** *Row with spontaneous fMRI: a-d* - MSE reconstruction and PMAT24 CR for EDR binary, EDR continuous, Geometry and EDR+LR respectively. *Row with task-evoked fMRI of mean 7 canonical tasks: a-d* - MSE reconstruction and PMAT24 CR for EDR binary, EDR continuous, Geometry and EDR+LR respectively.

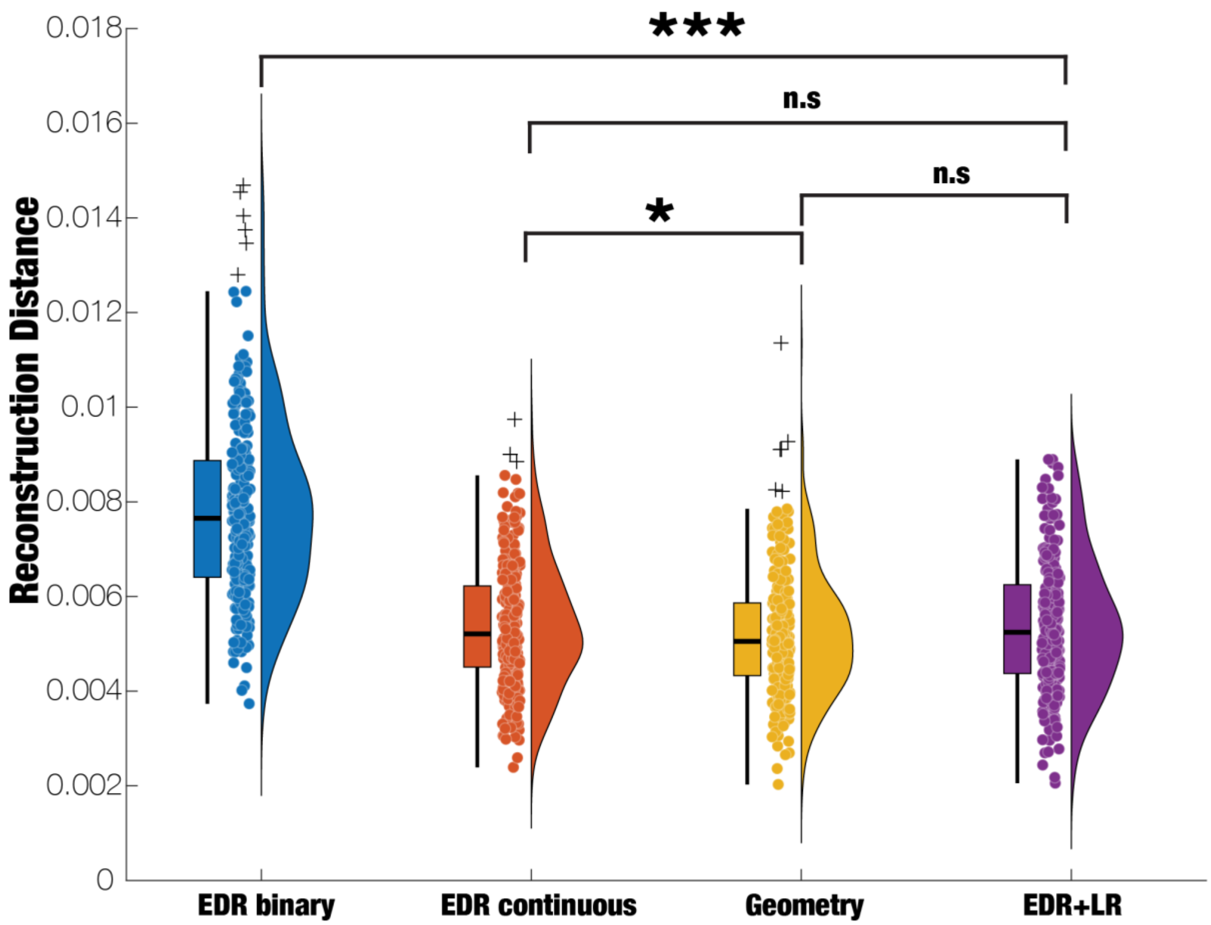


**Fig. S9. Applying global signal regression makes the reconstruction of EDR+LR and Geometry statistically non-significant.** The reconstruction of FC long-range connections with global signal regression (defined by high correlation values, >0.5 correlation, and Euclidean distance, <40mm) for 200 modes using all four graph representations. EDR+LR has non-significant MSE reconstruction with Geometry and EDR continuous (EDR+LR vs. EDR binary $p<10^{-3}$, EDR+LR vs. EDR continuous n.s., EDR+LR vs. Geometry n.s., and EDR continuous vs. Geometry p<0.0205, Bonferroni corrected two-tailed paired t-test, * p<0.05, ** p<0.01, *** p<0.001).

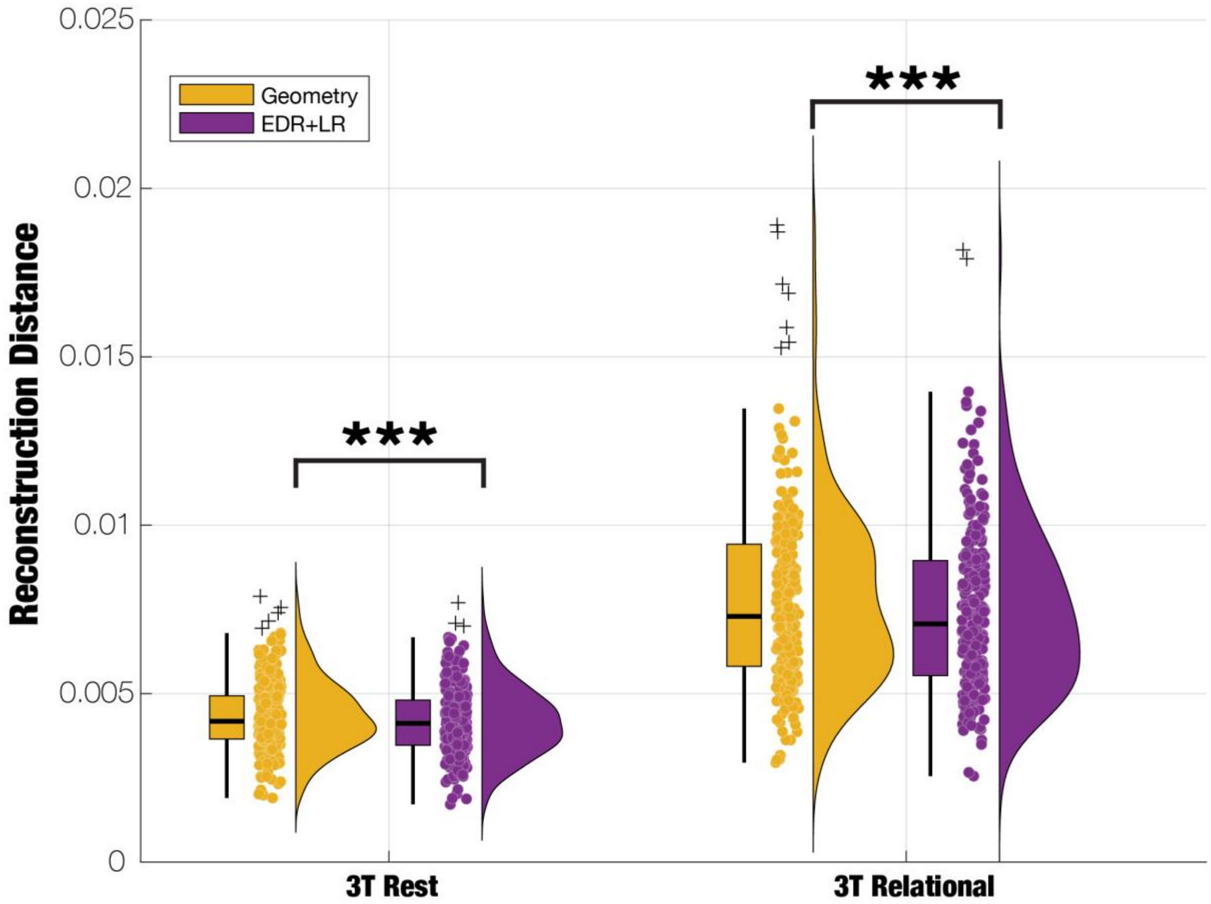


**Fig. S10. EDR+LR reconstruction of brain dynamics for spontaneous and relational tasks is significantly better than with Geometric reconstruction.** The reconstruction of FC long-range connections for 200 modes using both the Geometric and EDR+LR graph representations. EDR+LR has significantly lower MSE reconstruction for both spontaneous and relational task activity (3T Rest: $p<10^{-3}$ and 3T Relational: $p<10^{-3}$, two-tailed paired t-test, * $p<0.05$, ** $p<0.01$, *** $p<0.001$).

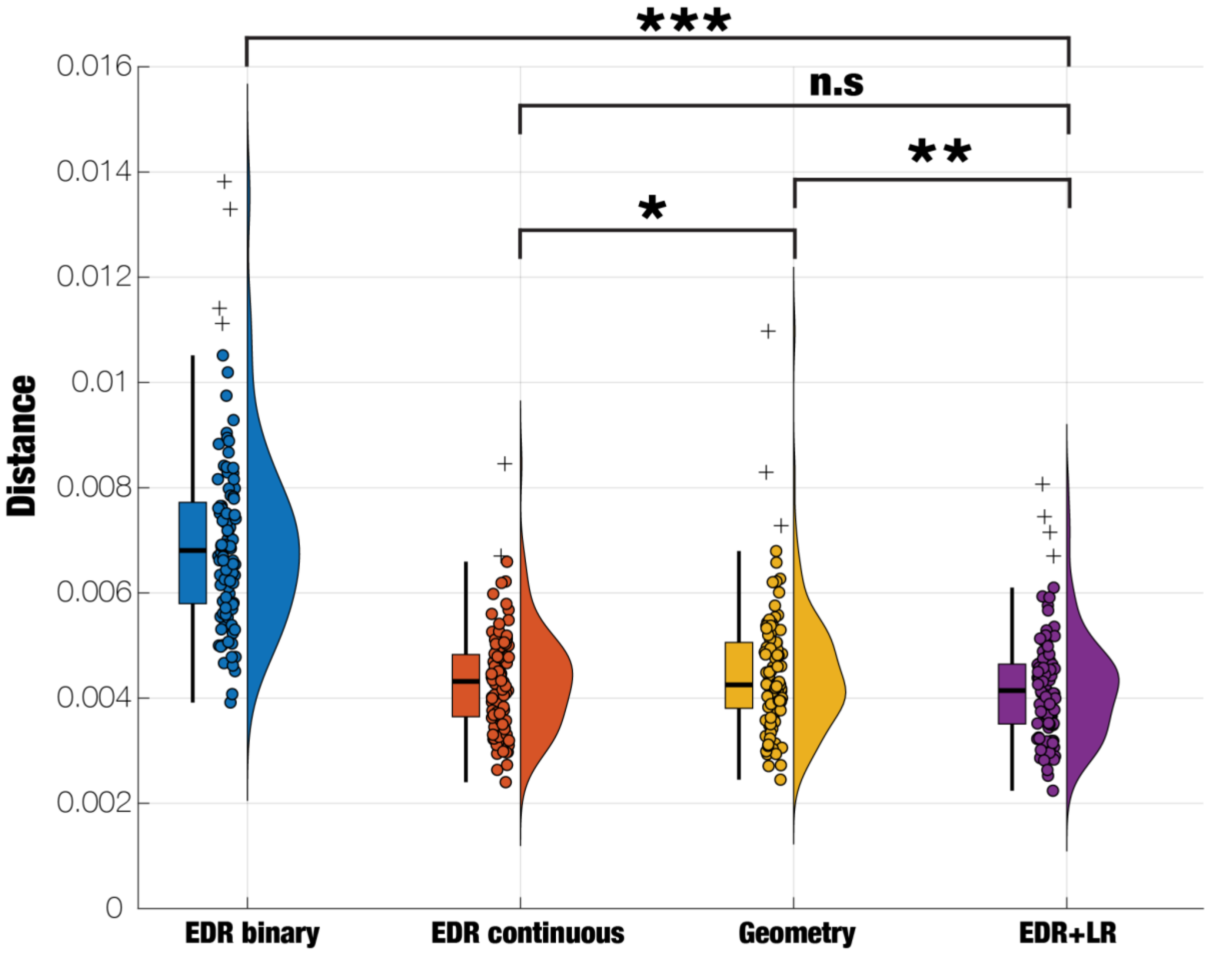


**Fig. S11. Reproducible findings of the main results on the reconstructions of long-range functional connectivity of spontaneous brain activity with an independent HCP cohort of 100 participants.** The reconstruction of FC long-range connections (defined by high correlation values, >0.5 correlation, and geodesic distance, >40mm) for 200 modes using all four graph representations. The results are consistent with the main analysis of reconstructing the spontaneous fMRI with the four graph representations (EDR+LR and Geometry $p<0.0023$, EDR+LR and EDR continuous n.s., EDR+LR and EDR binary $p<10^{-3}$, EDR continuous and Geometry $p<0.0356$, Bonferroni corrected two-tailed paired t-test, * $p<0.05$, ** $p<0.01$, *** $p<0.001$).
